# NF90/ILF3 is a transcription factor that promotes proliferation over differentiation by hierarchical regulation in K562 erythroleukemia cells

**DOI:** 10.1101/189647

**Authors:** Ting-Hsuan Wu, Lingfang Shi, Jessika Adrian, Minyi Shi, Ramesh V. Nair, Michael P. Snyder, Peter N. Kao

## Abstract

NF90 and splice variant NF110 are DNA‐ and RNA-binding proteins encoded by the Interleukin enhancer-binding factor 3 (*ILF3*) gene that regulate RNA splicing, stabilization and export. The role of NF90 in regulating transcription as a DNA-binding protein has not been comprehensively characterized. Here, ENCODE ChIP-seq identified 9,081 genomic sites specifically bound by NF90/110 in K562 cells. One third of binding sites occurred at promoters of annotated genes. NF90/110 binding colocalized with chromatin marks associated with active promoters and strong enhancers. Comparison with 150 ENCODE ChIP-seq experiments revealed that NF90 clustered with transcription factors exhibiting preference for promoters over enhancers (*POLR2A*, *MYC*, *YY1*). Differential gene expression analysis following shRNA knockdown of NF90 in K562 cells revealed that NF90 directly activates transcription factors that drive growth and proliferation (*EGR1*, *MYC)*, while attenuating differentiation along erythroid lineage (*KLF1*). NF90/110 binds chromatin to hierarchically regulate transcription factors to promote proliferation and suppress differentiation.

## INTRODUCTION

Eukaryotic gene expression depends on tight and dynamic regulation of RNA transcription. Transcription of RNA occurs pervasively, and is critically modulated by nucleic acid-binding proteins at the levels of epigenetic control of chromatin landscape, transcription of the genome from DNA into RNA, and regulated stability of the resulting transcript (Kornberg 1999). Dynamic regulation of gene expression confers organismal diversity (Levine and Tjian 2003), cell-type complexity (Goldberg et al. 2007), and underlies the immediate early response of a cell upon activation by external stimuli (Jaenisch and Bird 2003).

Chromatin regulation is crucial for control of eukaryotic RNA transcription. Chromatin is packed into repeating units of nucleosomes (Kornberg 1977). Binding of transcription factors to DNA is regulated by chromatin remodelers that unwind nucleosomes to expose the accessibility of *cis*-regulatory elements (Berger 2007; Li et al. 2007).

Open chromatin regions may be transcribed. Significant portions of the genome give rise to RNA. The eukaryotic transcriptome extends beyond annotated protein-coding genes to include non-coding RNAs (ncRNAs) (Kapranov et al. 2007; Jacquier 2009). Functional regulatory elements, such as enhancers, are also transcribed to produce enhancer RNAs (eRNAs) (Ren 2010). Analysis of transcription initiation revealed that promoters and enhancers share a unified architecture of tightly spaced divergent transcription start site (TSS) pairs, and bi-directional production of transcript (Core et al. 2014).

Stability of the RNA product is regulated after transcription. At promoters for protein-coding genes, the sense transcript is typically stable, while the upstream antisense RNA (uaRNA) is susceptible to rapid degradation (Preker et al. 2008). A pair of stable transcripts may also be produced at bidirectional promoters if the divergent transcription start site pairs each respectively encode mRNA, or if the mRNA is accompanied by a long non-coding RNA (lncRNA) (Wei et al. 2011).

DNA‐ and RNA-binding proteins (DRBPs) interact with diverse classes of nucleic acids and regulate RNA transcription on multiple levels. RNA-binding proteins are often studied in regulation of protein translation (Burd and Dreyfuss 1994). However, recent studies that identified a class of dual DNA‐ and RNA-binding proteins show that they may play a key role in modulating gene expression (Cassiday and Maher 2002; Hudson and Ortlund 2014).

Nuclear Factor 90 (NF90) is a protein encoded by the Interleukin enhancer-binding factor 3 (*ILF3*) gene first cloned as part of a multi-protein complex purified by DNA affinity chromatography from activated Jurkat T cell nuclear extract based on inducible and specific binding to the <u>N</u>uclear <u>F</u>actor of <u>A</u>ctivated <u>T</u> cell (NF-AT) target sequence in the IL-2 promoter (Corthesy and Kao 1994). The proteins that formed tight heterodimer comprising the inducible NF-AT binding complex was identified to be Nuclear Factor 90 (NF90, *ILF3*) and Nuclear Factor 45 (NF45, *ILF2*) (Kao et al. 1994).

Subsequent to their initial discovery based on inducible binding to the NF-AT sequence *in vitro*, NF90 and NF45 have been widely studied in diverse cellular processes such as DNA-break repair (Shamanna et al. 2011), cell cycle regulation (Shi et al. 2005; Guan et al. 2008; Jiang et al. 2015; Ni et al. 2015), cell growth and proliferation (Fung et al. 2000; Reichman et al. 2002; Rhodes et al. 2004; Ben-Porath et al. 2008; Huang et al. 2014; Ni et al. 2015; Higuchi et al. 2016; Jiang et al. 2017). In particular, NF90 is recognized as a double-stranded RNA binding protein that participates in regulation of translation by binding to post-transcriptional mRNA (Shim et al. 2002; Pei et al. 2008; Vumbaca et al. 2008; Zhu et al. 2010), microRNA processing (Sakamoto et al. 2009; Masuda et al. 2013), and identified as a host factor in viral replication (Isken et al. 2007; Wang et al. 2009; Gomila et al. 2011; Shabman et al. 2011; Li et al. 2014; Li and Belshan 2016).

NF90 has been extensively studied as a RNA-binding protein. However, the original purification using DNA affinity chromatography and subsequent ChIP experiments suggest a role for NF90 in gene regulation through its capacity to bind DNA. Here, we used chromatin immunoprecipitation of NF90 followed by deep sequencing (ChIP-seq) and integrated available ENCODE data with our ChIP-seq data in K562 cells to characterize its transcriptional regulation of genomic targets. We reveal a novel role for NF90 as a dual DNA‐ and RNA-binding protein and important regulator of chromatin, RNA transcription, and transcript stability.

## RESULTS

### NF90 binds the genome extensively

To analyze genome-wide binding sites of NF90, we used chromatin immunoprecipitation followed by deep sequencing (ChIP-seq) in the chronic myelogenous leukemic (K562) cell line, a ‘Tier 1’ cell line prioritized by the Encyclopedia of DNA Elements (ENCODE) Project. Abundant data mapped onto the hg19 (Human Genome version 19) reference genome from K562 experiments deposited by various ENCODE-affiliated groups allowed us to integrate our ChIP-seq analysis with studies on chromatin landscape, transcription factor binding, and gene expression analysis of the K562 genome to understand the role of NF90 in regulating gene expression in the context of diverse combinatorial events on the chromatin.

ChIP-seq revealed extensive occupancy of NF90 on the genome. Over 9,000 recovered binding peaks were consistent across Irreproducible Discovery Rate (IDR) analysis of biological replicates (Fig. 1a). We observed occupancy near annotated transcription start sites (TSS) (Fig. 1c), with over a third of binding peaks occurring at upstream promoter regions (Fig. 1b). A plurality of peaks occurred in distal intergenic regions (Fig. 1b). Examining the average binding profile of NF90 occupancy near all TSS, we found a conserved pattern featuring bimodal binding of NF90 with peaks flanking the TSS, with a stronger peak typically occurring downstream of the TSS. Along the entire transcribed loci of gene bodies, there is enrichment both near the TSS, and at the transcription end site (TES) (Fig. 1c).

**Figure 1.**
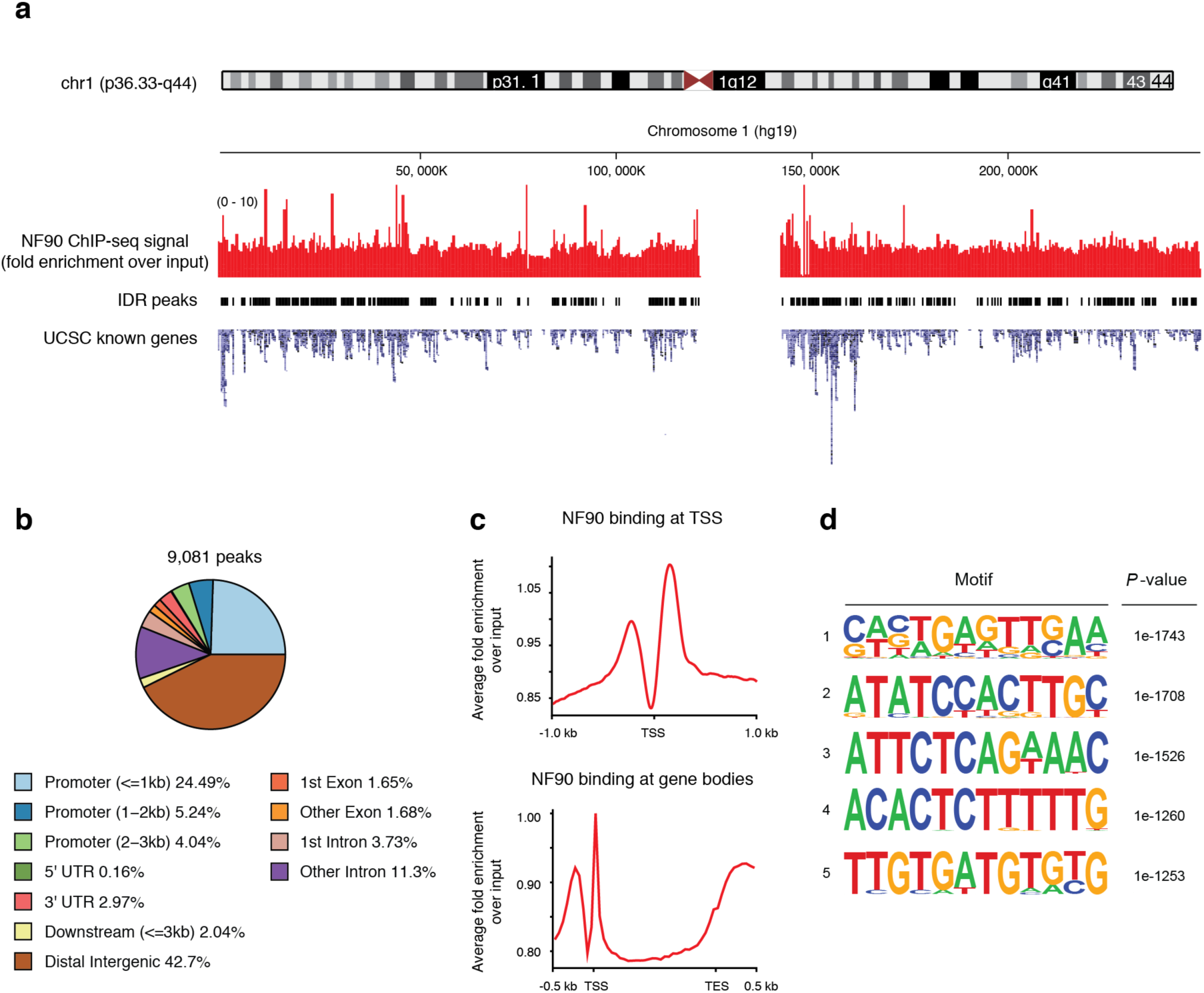
ChIP-seq reveals extensive NF90 occupancy along the genome. **(a)** ChIP-seq signal (fold enrichment over input) of NF90 shown along chromosome one, along peaks called using Irreproducibility Discovery Rate (IDR) between biological replicates. UCSC known genes and transcripts are aligned, **(b)** Distribution of annotated features nearNF90 binding sites, **(c)** Average binding profile of NF90 computed using all annotated transcription start sites (TSS). For gene bodies, all genes were scaled to a 1.5 kb length; the 0.5 kb flanking regions are not scaled, **(d)** Motif analysis (HOMER) based on NF90 ChIP-seq results in K562 cells showing enriched motifs in NF90 peaks. Shown: motifs ranked top 5 by statistical significance.

### NF90 binding sites are enriched for the NF-AT sequence

*De novo* motif discovery was performed on NF90 IDR peaks, and we recovered several motifs that were statistically enriched in the NF90 bound genomic regions (Fig. 1d). Notably, the 5’ – CTCTTTTT – 3’ (reverse complement: 5’ – AAAAAGAG – 3’) motif was discovered to be highly enriched (*P* = 1e-1260), which bore striking similarity to the original Nuclear Factor of Activated T cell (NF-AT)/ antigen receptor response element 2 (ARRE-2) target sequence in the Interleukin 2 (IL-2) promoter (5’ – GAGGAAAAAC – 3’). This result is consistent with original purification of NF90 from activated T cell nucleus using DNA affinity chromatography (Corthesy and Kao 1994; Kao et al. 1994), as well as subsequence chromatin immunoprecipitation studies that revealed dynamic binding of NF90 to the IL-2 promoter (Shi et al. 2005; Shi et al. 2007).

### NF90 binding is enriched at active promoters and enhancers

The exact binding pattern of a transcription factor depends on genomic accessibility controlled by dynamic chromatin landscape. We sought to study interplay between NF90 binding and different chromatin regions bearing distinct epigenetic marks.

Previously, Ernst and Kellis demonstrated that chromatin landscape of nine human cell types could be characterized into 15 distinct states by analyzing 14 genome-wide chromatin tracks using a hidden Markov model (Ernst et al. 2011). Several histone modification marks were used, as well as RNA polymerase II, and *CTCF*, a sequence-specific insulator protein.

Chromatin state were annotated into six broad classes, including: promoter, enhancer, insulator, transcribed, repressed, and inactive states. To understand the distributions of NF90 binding affinities for each chromatin state, over 9,000 NF90 ChIP-seq peaks were sorted based on their underlying chromatin state annotation.

We found significant NF90 binding in 8 of the 15 chromatin states, detailed in Figure 2. Observing all 9,081 NF90 ChIP-seq peaks confirmed by IDR analysis, we find most binding occurring in state 1 – active promoter (4,735 peaks), followed by state 4 – strong enhancer (2,479 peaks). Of all active promoter (State 1) regions annotated for the K562 genome, 30% were bound by NF90 (Fig. 2). The preference of NF90 occupancy at active promoters (State 1) and strong enhancers (State 4) based on chromatin state reinforces our finding that NF90 binding sites frequently occur near TSS (Fig. 1b). Moreover, NF90-bound regions are enriched for high levels of transcription.

**Figure 2.**
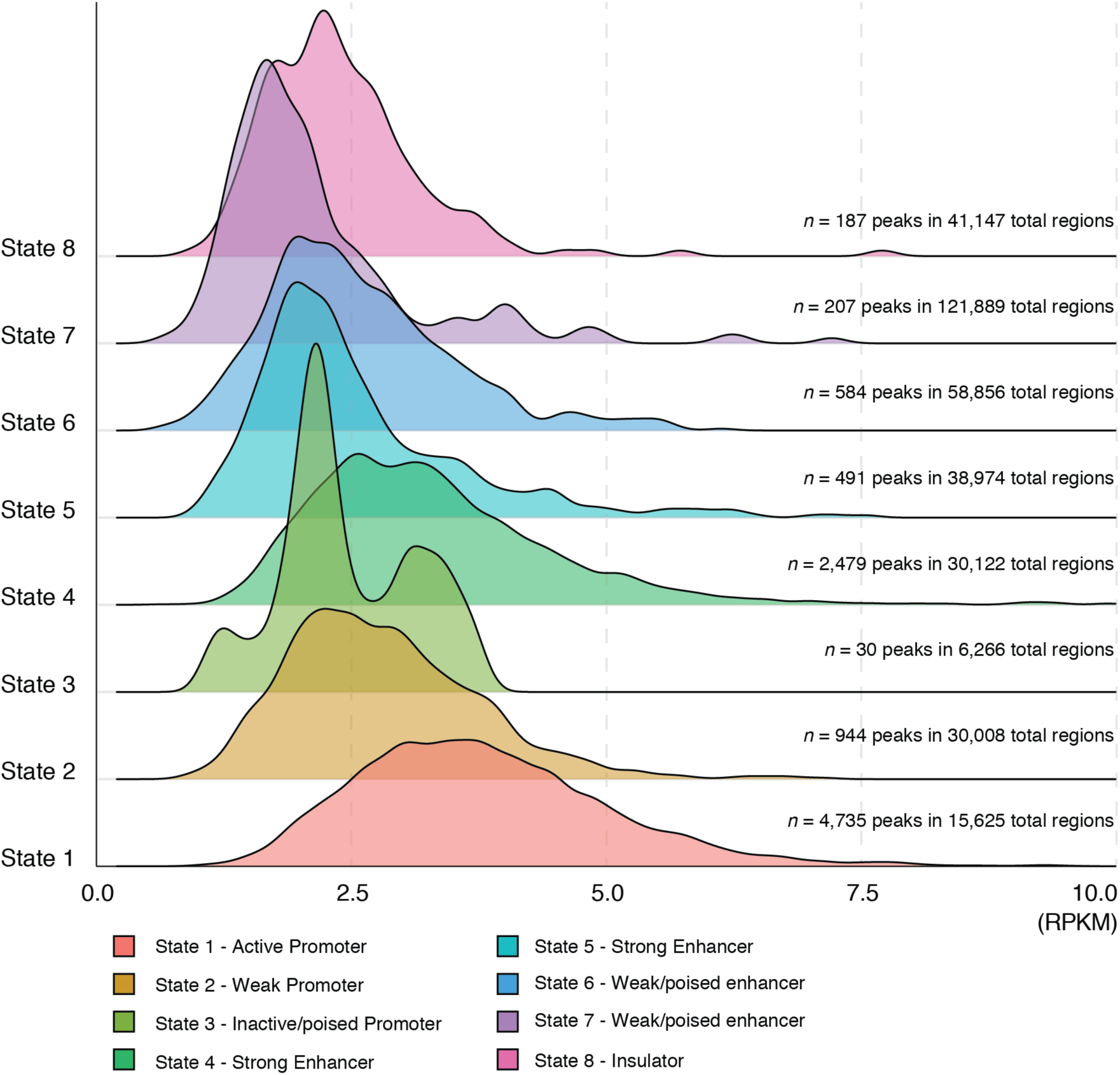
NF90 relative enrichment in different chromatin states. IDR peaks called from NF90 ChIP-seq experiment in K562 were sorted according to the chromatin state they resided in. The segmented peaks for each of 15 chromatin states were then used to query the ChIP-seq read files to count the number of reads to obtain affinity information for each peak. The resulting distribution of NF90 binding affinities in different chromatin states were plotted as a histogram, *x*-axis: 8 of 15 chromatin states in which NF90 peaks resided in. *y*-axis: Reads Per Kilobase of transcript per Million mapped reads (RPKM).

NF90 occupied active promoters and enhancers with higher frequency and tighter affinity compared to other chromatin states. NF90 peaks in active promoters (State 1) exhibited the highest binding affinities, followed by those in strong enhancers (State 4), and weak promoters (State 2). The normal shape of the histogram in these open chromatin regions indicates a continuous range of binding affinities of NF90, suggesting nonspecific and promiscuous association of the protein at these permissive chromatin states (Nie et al. 2012). This is consistent with previous studies that found motif depletion and nonspecific binding in permissive chromatin regions (Ernst and Kellis 2013).

### NF90 binding colocalizes with selective histone marks at active promoters and enhancers

Combinatorial histone marks partitioned the K562 genome by transcriptional activity (Ernst et al. 2011; Ernst and Kellis 2013), and may define distinct chromatin states (Calo and Wysocka 2013). To investigate if NF90 physically colocalized with the key histone marks that respectively defined promoter and enhancer states, we accessed histone mark ChIP-seq experiments performed on K562 deposited to ENCODE. The average binding profiles of NF90 alongside key histone marks were graphed at selected chromatin states.

We aligned NF90 ChIP-seq binding with the histone marks enriched for active promoter binding: RNA pol II, H3K9ac, and H3K4me3(Fig. 3a-c). Histone modifications H3K4me3 and H3K9ac both showed substantial binding, as expected; NF90 binding strength was comparable to RNA pol II at transcription start sites (Fig. 3a). The shapes of binding curves of H3K9ac and NF90 were similar, as both exhibited bimodal binding (Fig. 3a). At active promoters, the average binding profile of NF90, H3K4me3, H3K9ac, and RNA pol II showed remarkable colocalization of these peaks with mirrored binding strengths. All tracks consistently peaked at around 1000 bp downstream of defined center of active promoter regions (Fig. 3b). In contrast, when we examined average binding profiles at inactive promoters, we found diminished and noisy binding of NF90, along with H3K9ac and RNA pol II (Fig. 3c). At inactive promoters, only H3K4me3 still exhibited a discernible peak, indicating a broader association both with active and inactive promoters (Fig. 3c), consistent with previous literature (Creyghton et al. 2010).

**Figure 3.**
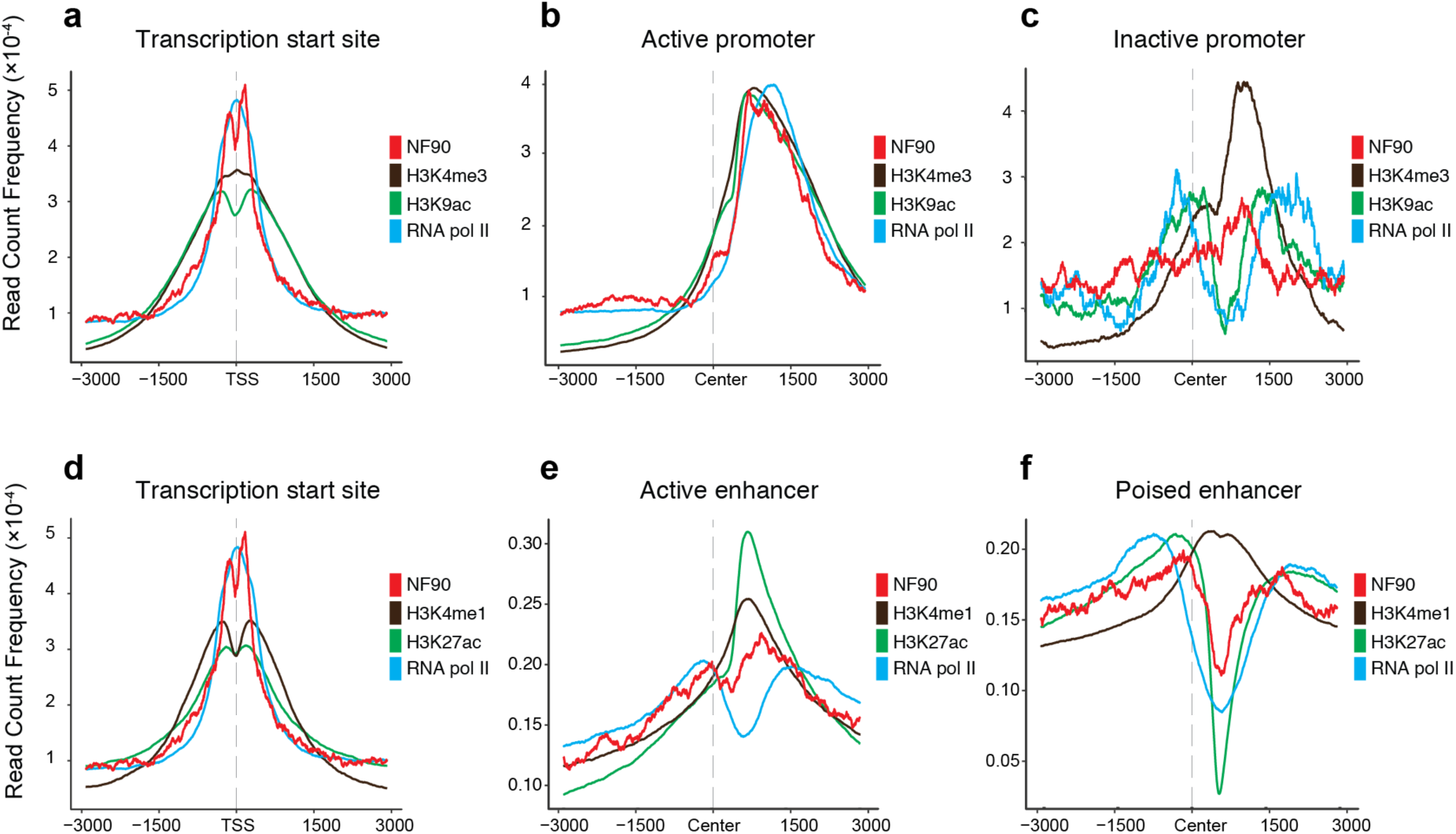
NF90 colocalization with specific histone marks at active promoters and enhancers. **a-c.** Average binding profile of NF90, H3K4me3, H3K9ac, and RNA pol II at **(a)** transcription start sites, **(b)** active promoters, and **(c)** inactive promoters, **d-f.** Average binding profile of NF90, H3K4mel and H3K27ac at **(a)** transcription start sites, **(b)** active enhancers, and **(c)** poised enhancers. Transcription start sites retrieved from UCSC Known Genes database. Active promoters, inactive promoters, active enhancers and poised enhancers from Ernst *et al*. *x*-axis: relative position near TSS. *y*-axis: read count frequency of tag within region.

Next, we aligned NF90 ChIP-seq binding with the histone marks enriched for active enhancer binding: RNA pol II, H3K27ac, and H3K4me1 (Fig. 3d-f). NF90 colocalized with H3K27ac and H3K4me1 at transcription start sites, and these histone modifications exhibited bi-modal binding with peaks that colocalized with NF90 peaks (Fig. 3d). At active enhancer regions, NF90 colocalized with histone marks H3K27ac and H3K4me1, with H3K27ac showing strongest binding, and NF90 showing binding strength comparable to H3K4me1 (Fig. 3e). All tracks consistently peaked at nearly 1000 bp downstream of center of enhancer region. In contrast, at poised enhancers, NF90 binding mirrored H3K27ac and both tracks exhibited a striking decrease at 1000 bp downstream of defined enhancer center. H3K4me1 still showed binding at poised enhancers with a defined peak (Fig. 3f), demonstrating more promiscuous binding, consistent with literature describing broader association of H3K4me1 with both active and poised enhancers (Creyghton et al. 2010).

### NF90 binding pattern by chromatin state clusters with sequence-specific transcription factors

Open chromatin is generally permissive to transcription factor binding. However, distinct classes of regulatory factors may show different preferences for different types of open chromatin (Ernst and Kellis 2013).

To investigate if the chromatin state preference of NF90 may cluster with similar regulatory factors, we accessed 150 datasets of transcription factor (TF) ChIP-seq peaks performed in K562 that were generated by the ENCODE analysis working group using a uniform processing pipeline. We computed the relative enrichment of all 150 TF peaks as well as NF90 ChIP-seq. An unsupervised *k*-means algorithm was used to cluster regulatory factors into four distinct classes.

Principal component analysis (PCA) revealed that the four classes are well separated along the first two principal components, indicating that much of the variance is explained by a relative enrichment along a few of the chromatin states (Fig. 4). The major eigenvectors plotted show that most variance among regulatory factors in the original dataset is explained by the relative enrichment in: State 1 – active promoter, State 4 – strong enhancer, and State 8 – insulator (Fig. 4)

**Figure 4.**
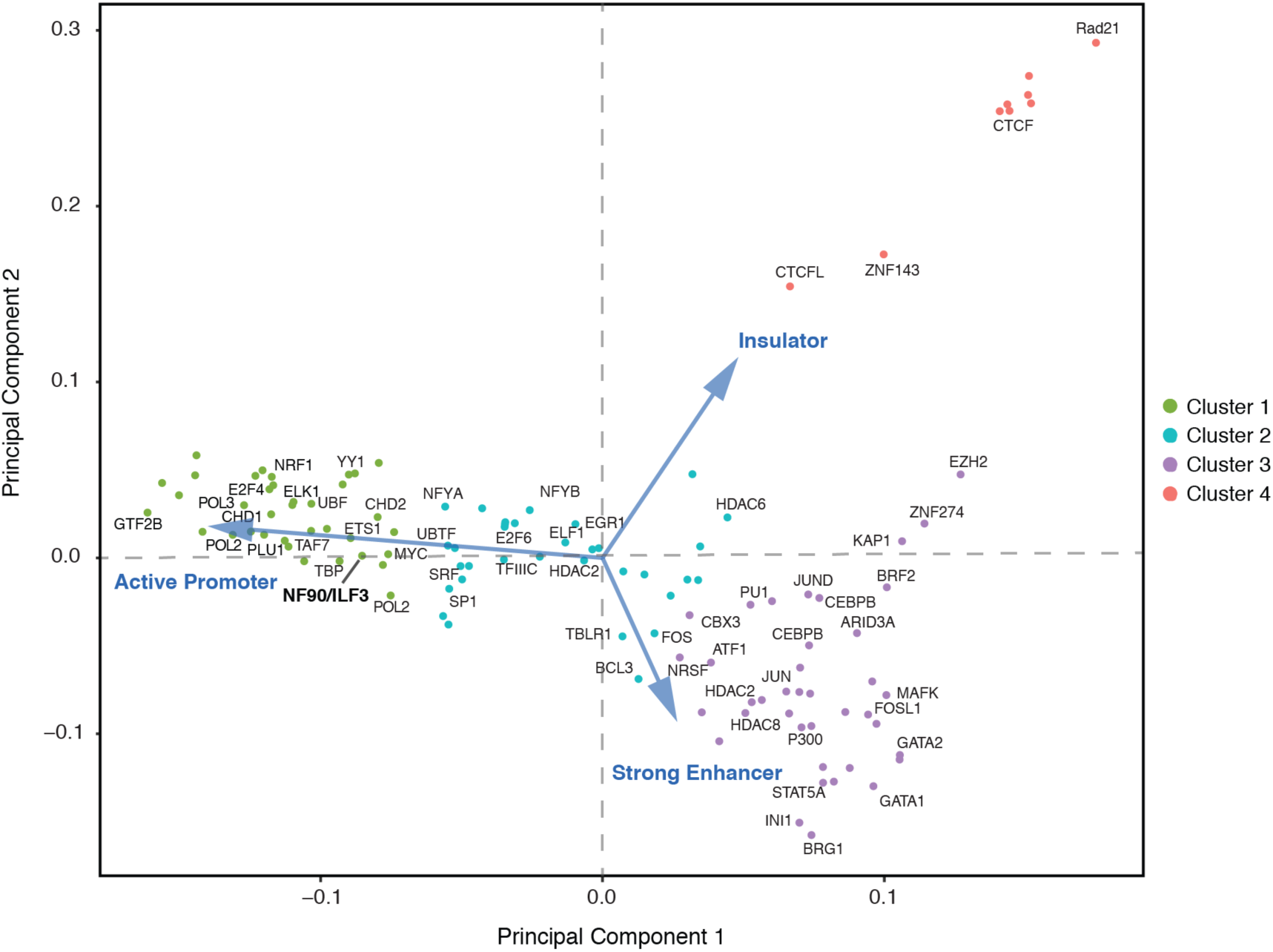
Clustering of transcription factors based on relative enrichment for chromatin states. 150 datasets of transcription factor ChIP-seq peaks based on ENCODE data production centers processed through a uniform processing pipeline were retrieved. 119 unique regulatory factors, including generic and sequence-specific factors were retrieved for the K562 line to supplement the NF90 ChIP-seq data. For a given transcription factor ChIP-seq peak set, the relative enrichment in different chromatin states was computed. Enrichments were then row-normalized by the largest enrichment values for each experiment. *k*-means clustering with *K* = 4 produced the clusters graphed here using principal component analysis (PCA). The major eigenvectors for the original dataset is depicted in blue arrows.

NF90 binding was strongly biased towards active promoters, consistent with our findings before (Fig. 2). NF90 chromatin occupancy most resembled other sequence-specific transcription factors in Clusters 1. Notably, NF90 clusters closely with the proto-oncogenic transcription factors *MYC* and *ETS1*. In contrast, the regulatory factors in Cluster 3 with preference for strong enhancers included the histone acetyltransferase *EP300*, which may represent general coactivator that do not bear strong sequence-specific binding to the genome. NF90 is well separated from the regulators in Cluster 4, primarily *CTCF* insulating proteins with transcriptional repression activity.

### NF90 binding at divergent transcription initiation regions is enriched for transcript stability

A substantial fraction of the eukaryotic genome is transcribed, with bidirectional initiation resulting in transcripts of varying stability. Core *et al*. previously characterized all pairs of TSS found along the K562 genome by measuring *de novo* transcription using a variation of Global Run-On Sequencing that enriches for 5’-capped RNAs (GRO-cap). The GRO-cap signal is selective for transcription initiation events, while sustained GRO-seq tends to detect stable transcripts. Thus, the stability of transcripts at each TSS pair could be characterized as stable or unstable using a hidden Markov model (Core et al. 2014).

To investigate if NF90 binding is persistent at TSS pairs that define divergent transcription initiation, we retrieved the coordinates for all *de novo* TSS pairs in the K562 genome discovered using GRO-cap. The average binding profile of NF90 at *de novo* TSS pairs was bimodal with peaks flanking the center of the region (Fig. 5a), resembling what we found at annotated TSS (Fig. 1c).

**Figure 5.**
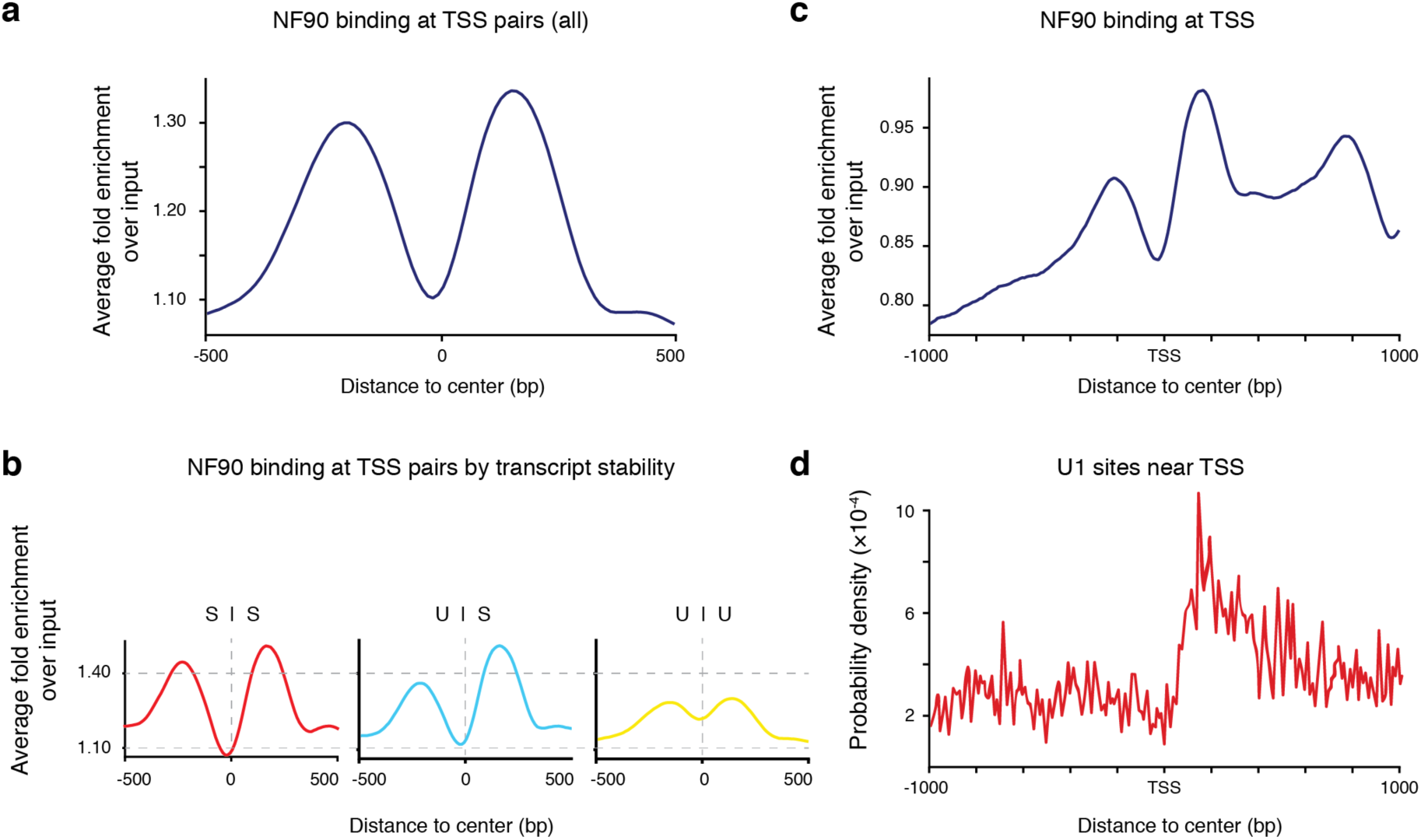
NF90 binding at divergent transcription initiation regions is enriched for transcript stability. **(a)** Average binding profile of NF90 ChIP-seq peaks at all transcription start site (TSS) pairs, **(b)** Average binding profile of NF90 ‐seq peaks at transcription start site (TSS) pairs sorted by transcript stability parity. SS = stable – stable, US = unstable – stable, UU = unstable – unstable, **(c)** Average binding profile of NF90 ChIP-seq peaks centered at TSS with lkb flanking regions, **(d)** Local motif enrichment analysis of the U1 SS5 splice sequence centered at TSS with lkb flanking regions. TSS pair coordinates from Core *et al*.

Bimodal binding of NF90 at TSS pairs suggests selective association with individual transcripts rather than shared machinery. To investigate whether NF90 binding affinity at individual transcripts correlated with stability, we subsetted the original collection of TSS pairs into: stable-stable, unstable-stable, and unstable-unstable transcript pairs, as originally described by Core *et al*. NF90 bound stable-stable TSS pairs equally well, replicating the bimodal enrichment found in all TSS pairs (Fig. 5b). Strikingly, the NF90 binding profile at unstable-stable TSS pairs demonstrated preferential enrichment at stable transcripts, and at unstable-unstable TSS pairs there was equal depletion of NF90 binding (Fig. 5b).

DNA sequence is known to influence the stability of the resulting RNA transcript (Bregman et al. 2011). The U1 splicing complex directed by the U1 small nuclear ribonucleoprotein (snRNP) to 5’ splice sites (SS5) may suppress polyadenylation-dependent termination, protecting the stability of the nascent transcript through productive elongation. Previously, NF90 was shown to interact with multiple components of the spliceosome (Chaumet et al. 2013). We investigated if NF90 binding preference at stable transcripts was related to presence of the U1 SS5 motif. All TSS were searched for enrichment of the U1 SS5 motif, and this enrichment was compared to the relative occupancy of NF90 near TSS. We found a striking colocalization of NF90 peak binding with an enrichment of the U1 SS5 site around 200 bp downstream of TSS (Fig. 5c-d). These observations suggest that NF90 may participate in regulating transcript stability through initial recruitment to the U1 SS5 site on the DNA and subsequent involvement in splicing machinery with the RNA transcript.

### NF90 regulates transcription of its genomic targets

To investigate the functional role of NF90 in transcriptional regulation, we accessed ENCODE RNA-seq data in K562 cells treated with shRNA directed towards NF90. We accessed this ENCODE published dataset to measure the global effects on the transcriptome of K562 cells upon NF90 knockdown, and intersected these differentially expressed genes with our NF90 ChIP-seq dataset to identify genes under direct transcriptional regulation by NF90.

Knockdown of NF90 resulted in 446 genes to be differentially expressed. The posterior probability of differential expression (PPDE) versus log2-transformed Fold Change (FC) was graphed for each gene (Fig. 6a). There was no obvious preference in directionality of change in gene expression upon NF90 knockdown; about 20% more genes were down-regulated (246 genes) compared to up-regulation (200 genes) (Fig. 6a). A list of 2,277 genes for which NF90 occupancy was discovered in its proximal promoter region was retrieved. These 2,277 genes were then compared to the 466 differentially expressed gene upon NF90 knockdown to identify genes that are directly transcriptionally regulated by NF90 (Fig. 6b). The resulting intersection of the NF90-bound genes with the differentially expressed genes is statistically significant (Fisher’s exact test, *P* = 1.1 e-04) with integrated set of 89 genes under direct regulation of NF90 (Fig. 6b).

**Figure 6.**
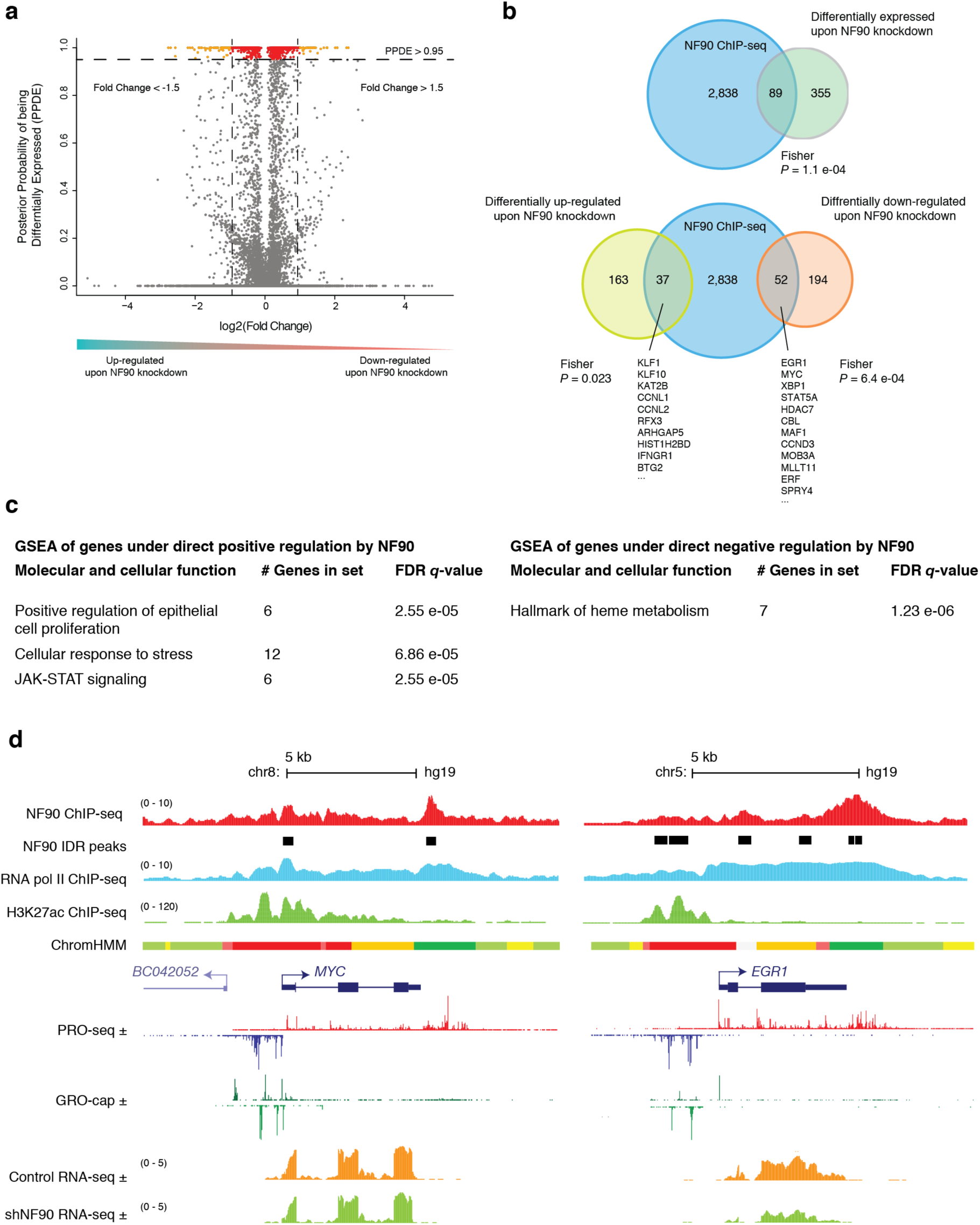
NF90 regulates transcription of its genomic targets. **(a)** Volcano plot demonstrating statistically significant changes in gene expression between control and NF90 knockdown in K562. *x*-axis: log fold change in shNF90 compared to control. *y*-axis: posterior probability of being differentially expressed (PPDE). Each dot is a gene; red: PPDE > 0.95; orange: PPDE > 0.95 and fold change > 1.5. **(b)** Venn diagram analyzing overlap between NF90 ChIP-seq genic targets and differentially expressed gene upon NF90 knockdown in K562 (*P* = 1.4 e-08). Venn diagram analyzing overlap between NF90 ChIP-seq genic targets, up-regulated genes upon NF90 knockdown, and down-regulated genes upon NF90 knockdown. **(c)** Gene set enrichment analysis (GSEA) of genes under direct positive or negative regulation by NF90. **(d)** Representative ChIP-seq track alignment of NF90, RNA pol II, H3K27ac; Chromatin state annotations from ChromHMM (red: active promoter, light red: weak promoter, orange: active enhancer, yellow: weak enhancer, dark green: transcriptional transition/elongation, light green: weakly transcribed); Precision nuclear run-on sequencing (PRO-Seq) and GRO-cap reads in reads per million (RPM); RNA-seq unique reads track alignment of control shRNA, and shNF90.

We further interrogated the effect of NF90 transcriptional regulation by separately intersecting the differentially down-regulated genes or the up-regulated upon NF90 knockdown, with the NF90 ChIP-seq promoter peaks. Out of 200 genes that were differentially up-regulated upon NF90 knockdown, NF90 binding was found in the promoter of 37 genes (Fig. 6b, Fisher’s exact test, *P* = 0.023). These genes are under direct negative regulation by NF90. Out of 246 genes that were differentially down-regulated upon NF90 knockdown, NF90 binding was found in the promoter of 52 genes (Fig. 6b, Fisher’s exact test, *P* = 6.4 e-04). These genes are under direct positive regulation by NF90.

Gene set enrichment analysis (GSEA) was used to identify sub-classes of genes associated with specific pathways that are over-represented in a gene set, and revealed the functional profile of genes under direct regulation by NF90. The 37 genes under direct negative regulation of NF90 was significantly enriched for Hallmark of heme metabolism (Fig. 6c) and differentiation down the erythroid lineage. Among which, Krueppel-like factor 1 (*KLF1*), a transcription factor with critical function in specifying hematopoietic differentiation and required for erythroid maturation, is bound by NF90 ChIP-seq analysis and differentially up-regulated upon knockdown (PPDE = 1, FC = 0.67).

GSEA of genes under direct positive regulation of NF90 revealed statistically significant overrepresentation of proto-oncogenes involved in Positive regulation of Epithelial cell proliferation (Fig. 6c). Moreover, genes involved in Cellular response to stress, and JAK-STAT signaling (Fig. 6c) indicate that NF90 is involved in the rapid transcriptional response upon extracellular signaling, consistent with its described role in regulating inducible expression of IL-2 upon T cell stimulation (Shi et al. 2005; Shi et al. 2007), as well as regulation of *FOS* immediate early expression upon stimulation (Nakadai et al. 2015).

A specific example of NF90 binding at the *MYC* locus is shown in Figure 6d. Divergent transcription can be observed wherein the upstream antisense RNA (uaRNA) to *MYC* is transcribed initially, as assayed by strong GRO-cap signal observable on both plus and minus strands (Fig. 6d). However, in contrast to the stable transcription observed at the *MYC*-coding plus strand as sustained PRO-seq signal across the entire locus, the minus strand PRO-seq signal for the uaRNA diminishes (Fig. 6d). At this stable-unstable TSS pair, NF90 signal is not observed along the uaRNA, but is enriched for the *MYC* TSS, and indeed along the entire transcribed *MYC* locus. Upon NF90 knockdown, transcription of *MYC* is significantly depleted, though fold change is modest. (PPDE = 1, FC = 1.67). Early growth response 1 (*EGR1*) is also identified as a target that is under direct positive regulation by NF90, and a similar phenomenon of NF90 binding along the body of the transcribed gene is illustrated at the *EGR1* locus (Fig. 6d), for which transcription is also significantly depleted upon NF90 knockdown (PPDE = 1, FC = 5.24).

Thus, inspecting the genes under direct transcriptional regulation by NF90 revealed that NF90 directly activates genes that drive growth and proliferation (*EGR1*, *MYC)*, while attenuating differentiation along the erythroid lineage (*KLF1*). NF90 interacts with chromatin at specific sites to hierarchically regulate transcription factors that promote cell proliferation and suppress differentiation.

## DISCUSSION

### NF90 as a hierarchical regulator of pluripotency and differentiation

Though previous studies mainly focused on the regulation of posttranscriptional processes by NF90 and its RNA-binding capacity, our results identify a role for NF90 in the transcriptional regulation of transcription factors by interacting with chromatin at specific sites, pointing to a hierarchical role of NF90 in the regulatory network of gene transcription.

Previously, Ye and colleagues (Ye et al. 2017) screened RNA binding proteins necessary for maintenance of pluripotency in embryonic stem cells (ESCs). Targeted disruption of NF90 and NF45 impaired ESC proliferation and promoted differentiation down embryonic lineages to an epiblast-like state.

Our integrated analysis of genes under direct regulation by NF90 supported its role in maintaining pluripotency and suppressing differentiation. The over-representation of proto-oncogenes involved in epithelial cell proliferation under direct positive regulation of NF90 suggests a role for NF90 in regulating growth and proliferation. This is consistent with previous studies of NF90 involvement in oncogenesis by regulating cell growth, cell cycle, and proliferation (Jiang et al. 2015; Higuchi et al. 2016; Jiang et al. 2017). Conversely, genes under direct negative regulation by NF90 included Krueppel-like factor 1 (*KLF1*), a critical regulator of hematopoietic development and specifier of the mature phenotype of the erythroid lineage (Vinjamur et al. 2014).

These results suggest a role for NF90 and its heterodimeric partner NF45 as the hierarchical regulators of the push and pull of pluripotency and differentiation by transcriptional control of key transcription factors (Ye et al. 2017).

### Modulation of NF90 DNA-binding and transcriptional activity

DNA‐ and RNA-binding proteins (DRBPs) assume an integral role in control of gene expression through versatile interactions with diverse nucleic acids. Consistent with its original biochemical purification of NF90 using DNA-affinity chromatography (Corthesy and Kao 1994) and subsequent electrophoretic mobility shift assay (EMSA) that showed a sequence-specific DNA binding activity *in vitro* (Kao et al. 1994), our ChIP-seq results demonstrate that NF90 associates with DNA *in vivo*.

These results taken together with the body of work on NF90 as an RNA-binding protein show that NF90 is a dual DNA‐ and RNA-binding protein. NF90 has two dsRNA binding domains (dsRBDs), and two additional domains that may interact with nucleic acids, both located at the C-terminus: an arginine‐ and glycine-rich RGG motif, and a GQSY domain (Parrott et al. 2005; Cazanove et al. 2008). Still lacking is identification of a definitive DNA-binding domain that mediates the wide association between NF90 and the chromatin described here, and how the DNA‐ and RNA-binding affinities of NF90 are regulated.

Functional enhancer elements have recently been recognized to transcribe noncoding RNAs, termed enhancer RNAs (Li et al. 2016). In the CRISPR/Cas9 system, noncoding RNAs complementary to DNA target sequences have been shown to form nuclear ribonucleoprotein particles that localize enzymatic activity to specific loci in the genome (Jinek et al. 2012). Single stranded RNA may also invade double-stranded DNA forming RNA:DNA hybrid R-loops. Mapping of RNA:DNA hybrids across the human genome revealed extensive binding and colocalization with the H3K27ac/H3K4me1 DNAse hypersensitivity epigenetic signature of actively transcribed promoters (Nadel et al. 2015).

Furthermore there was motif enrichment for GGAA sequences in the RNA components of the RNA:DNA hybrids. Mass spectrometry identified NF45/ILF2 and NF90/ILF3 proteins to be specifically associated with the RNA:DNA hybrids. We propose that nuclear noncoding RNAs may bind to NF90 and direct its transcriptional activation activity to diverse sites in the genome through RNA:DNA hybridization. The transcription factor YY1 is recognized to bind DNA and RNA and was successfully recruited through tethered RNA to different enhancers in embryonic stem cells (Sigova et al. 2015).

### NF90 coordination of transcription and translation

NF90 may coordinate transcription and translation of target genes (Castella et al. 2015). Coordinated regulation of transcription and translation by NF90 on the same target gene has been reported before (Shim et al. 2002; Shi et al. 2005; Pei et al. 2008). NF90 participates in regulating inducible expression of IL-2 upon T cell activation on multiple levels. Upon T cell stimulation, NF90 associates at the IL-2 proximal promoter dynamically (Shi et al. 2007), binds to the 3’UTR of the transcribed IL-2 mRNA (Pei et al. 2008), and mediates nuclear export of mature IL-2 transcript to the cytoplasm through the interaction of NF90 nuclear export signal with exportin (Shim et al. 2002). NF90 regulates inducible expression of endothelial growth factor (VEGF) upon hypoxia by binding to the human VEGF 3' untranslated region on the transcript (Vumbaca et al. 2008). Examining our NF90 ChIP-seq data, NF90 binds to the VEGFA promoter.

At the *FOS* locus, NF90 and NF45 were previously found to associate dynamically along the gene body, correlating spatially and temporally with that of RNA pol II occupancy (Nakadai et al. 2015). Consistent with this study, our ChIP-seq results have shown that in addition to binding at active promoters and enhancers, NF90 is associated extensively along the gene bodies of *EGR1* and *MYC* (Fig. 6d), which were also identified to be targets of transcriptional regulation by NF90. We note that the transcriptional regulation and extensive intragenic association of NF90 raises an intriguing possibility that NF90 may bind to and control the post-transcriptional processing of the mRNA synthesized from the very genomic targets under transcriptional control by NF90.

Van Nostrand et al. previously developed enhanced UV crosslinking and immunoprecipitation (eCLIP) for transcriptome-wide discovery of RNA-binding protein binding targets (Van Nostrand et al. 2016). The transcriptome-wide interactions of NF90 were characterized by eCLIP-seq and deposited on ENCODE. NF90 associated with the transcriptome extensively, binding to over 10,000 transcripts. In nearly all genes where NF90 binding was detected on the genome by ChIP-seq, there was corresponding eCLIP signal indicating NF90 binding to the resulting transcript. This is consistent with previous studies that have found promiscuous association of NF90 with its RNA binding partners with no obvious sequence specificity (Parrott et al. 2007).

Thus, we propose that NF90 may participate in transcription and translation of targets by continued recruitment to enhancer and promoter regions on the chromatin; transfer from chromatin to transcriptional machinery to allow dynamic association along gene body, then association with nascent transcript. In this model, NF90 functions analogously to a conveyor belt from chromatin, gene, to transcript, in the dynamic and ongoing process of protein synthesis.

## MATERIALS AND METHODS

### Data Accession

The following datasets were downloaded from the ENCODE Data Coordination Center (DCC): H3K4me1 ChIP-seq on human K562 (ENCFF159VKJ), H3K4me3 on human K562 (ENCFF616DLO), H3K9ac ChIP-seq on human K562 (ENCFF306MNO), H3K27ac ChIP-seq on human K562 (ENCFF038DDS), POLR2A ChIP-seq on human K562 (ENCFF285MBX), RNA-seq on K562 cells treated with an shRNA knockdown against ILF3 (ENCFF845BGZ, ENCFF153BJQ), Control shRNA against no target in K562 cells followed by RNA-seq (ENCFF439FIP, ENCFF702YIW).

### Chromatin immunoprecipitation sequencing (ChIP-seq)

#### Data acquisition and sequencing

All analysis was performed on the Human genome build 19 (GRCh37/hg19) reference genome. ChIP-seq of NF90 in K562 was performed by the Snyder Data Production Center (DPC) as part of the ENCODE consortium. Antibody used was against NF90 (mAb DRBP76; BD). ChIP-seq experiment in K562 protocol, quality control, and preprocessing followed ENCODE standards as part of the ENCODE uniform processing pipeline. (Consortium 2012; Landt et al. 2012).

#### Retrieval of genomic coordinates

Genomic coordinates of active promoters, inactive promoters, active enhancers, and poised enhancers in the human genome assembly 19 (hg19) were retrieved from the UCSC Table Browser (http://rohsdb.cmb.usc.edu/GBshape/cgi-bin/hgTables), based on Chromatin state discovery and characterization (ChromHMM) annotation from ENCODE/Broad Institute using ENCODE ChIP-seq data for nine histone modifications in K562 cells performed by Ernst and Kellis (Ernst and Kellis 2013).

Genomic coordinates of transcription start site (TSS) pairs, including the stability classification for stable-stable, stable-unstable, or unstable-unstable transcripts, were retrieved from the UCSC Table Browser based on PRO-seq and GRO-cap analysis in K562 performed by Core *et al* (Core et al. 2014).

#### Data analysis

Genome-wide peak coverage analysis and average binding profile for NF90 ChIP-seq data was performed in R with the ChIPseeker package and DeepTools (Yu et al. 2015; Ramirez et al. 2016). Motif enrichment analysis and gene annotation of NF90 ChIP-seq peaks were computed using Hypergeometric Optimization of Motif EnRichment (HOMER) (Heinz et al. 2010). NF90 binding affinity in each chromatin state was computed in R with the DiffBind package (Stark and Brown 2012).

#### Relative enrichment and cluster analysis

Relative enrichment of each regulatory factor in different chromatin states were computed essentially as described by Ernst and Kellis (Ernst and Kellis 2013). Briefly, to compute the enrichment for a peak call-data set in a specific chromatin state, *s*, we computed the enrichment for transcription factor binding as (*a*_*s*_/*b*)/(*c*_*s*_/*d*), where *a*_*s*_ is the total number of bases in a peak call in *s*; *b* is the total number of bases in a peak call; *c*_*s*_ is the total number of bases in s; and *d* is the total number of bases for which the segmentation was defined.

For cluster analysis, a single vector of *n* = 15 relative enrichment values for each chromatin state was obtained for *m* = 150 regulatory factors, and concatenated to form a *n*×*m* matrix. Unsupervised learning was performed in R using the K-means function using a range of *K* values. The principal component analysis (PCA) of the clusters were plotted, and *K* = 4 was chosen based on PCA analysis.

## RNA-seq analysis

### Differential gene expression analysis (EBSeq)

Gene quantification data was accessed from ENCODE: RNA-seq on K562 cells treated with an shRNA knockdown against ILF3 (ENCFF845BGZ, ENCFF153BJQ), Control shRNA against no target in K562 cells followed by RNA-seq (ENCFF439FIP, ENCFF702YIW). Differential gene expression analysis was performed in R using EBSeq package (Leng et al. 2013), an empirical Bayes hierarchical model for inference in RNA-seq experiments, using false discovery rate of 0.05 to retrieve list of 446 differentially expressed genes upon NF90 knockdown.

### Gene set enrichment analysis

Differentially expressed genes were ranked by posterior fold change and enriched for gene sets using The Molecular Signatures Database (MSigDB) and Gene Set Enrichment Analysis (GSEA) tools developed by the Broad Institute.

## DATA ACCESS

ChIP-seq data have been deposited with GEO Accession GSE10325, and at ArrayExpress accession E-MTAB-6042

UCSC browser session thsuanwu_ILF3ChIP_20170807: http://genome.ucsc.edu/cgi-bin/hgTracks?hgS_doOtherUser=submit&hgS_otherUserName=peterkao&hgS_otherUserSessionName=thsuanwu_ILF3ChIP_20170807

## ACKNOWLEDGEMENTS

Supported in part by NIH R01-AI39624 to PNK and the Hudson Fund for Allergy Research

